# Unraveling Axonal Transcriptional Landscapes: Insights from iPSC-Derived Cortical Neurons and Implications for Motor Neuron Degeneration

**DOI:** 10.1101/2024.03.26.586780

**Authors:** Jishu Xu, Michaela Hörner, Maike Nagel, Perwin Perhat, Milena Korneck, Marvin Noß, Stefan Hauser, Ludger Schöls, Jakob Admard, Nicolas Casadei, Rebecca Schüle

## Abstract

Neuronal function and pathology are deeply influenced by the distinct molecular profiles of the axon and soma. Traditional studies have often overlooked these differences due to the technical challenges of compartment specific analysis. In this study, we employ a robust RNA-sequencing (RNA-seq) approach, using microfluidic devices, to generate high-quality axonal transcriptomes from iPSC-derived cortical neurons (CNs). We achieve high specificity of axonal fractions, ensuring sample purity without contamination. Comparative analysis revealed a unique and specific transcriptional landscape in axonal compartments, characterized by diverse transcript types, including protein-coding mRNAs, RNAs encoding ribosomal proteins (RPs), mitochondrial-encoded RNAs, and long non-coding RNAs (lncRNAs). Previous works have reported the existence of transcription factors (TFs) in the axon. Here, we detect a set of TFs specific to the axon and indicative of their active participation in transcriptional regulation. To investigate transcripts and pathways essential for central motor neuron (MN) degeneration and maintenance we analyzed *KIF1C-knockout (KO)* CNs, modeling hereditary spastic paraplegia (HSP), a disorder associated with prominent length-dependent degeneration of central MN axons. We found that several key factors crucial for survival and health were absent in *KIF1C-KO* axons, highlighting a possible role of these also in other neurodegenerative diseases. Taken together, this study underscores the utility of microfluidic devices in studying compartment-specific transcriptomics in human neuronal models and reveals complex molecular dynamics of axonal biology. The impact of *KIF1C* on the axonal transcriptome not only deepens our understanding of MN diseases but also presents a promising avenue for exploration of compartment specific disease mechanisms.

## Introduction

The inaccessibility of the brain makes analysis of homeostatic and disease conditions challenging. However, understanding of these conditions and their pathological mechanisms is a prerequisite for developing targeted therapies for diseases of the central nervous system (CNS). Even though considerable research goes into the development of novel therapeutic approaches for common and rare neurodegenerative diseases, many are still without cure. Up to this day, mostly post-mortem derived brain tissue is analyzed, where preservation of RNA, protein, DNA, and lipids is challenging. Therefore, incomplete understanding of pathomechanism contributes to missing therapeutic solutions. Neurons are highly polarized and display high morphological complexity. Potent and functional communication between their soma and distal processes (axons) is necessary for correct functions (Bentley & Banker, 2016). Consequently, correct localization of mRNA and subsequent local protein synthesis are needed (Sutton & Schuman, 2006; Wang *et al*., 2010; Perry & Fainzilber, 2014; Tom Dieck *et al*., 2014). Interestingly, in neurodegenerative conditions like amyotrophic lateral sclerosis (ALS), Alzheimer’s disease (AD), Huntington disease (HD), glaucoma, and hereditary spastic paraplegia (HSP), axonal degeneration often precedes and sometimes is causative of neuronal death (Li *et al*., 2001; Ferri *et al*., 2003; Fischer *et al*., 2004; Libby *et al*., 2005; Stokin *et al*., 2005; Hörner *et al*., 2022). Additionally, recent studies provide evidence that mRNAs are present in large quantities in the axon, implicating local translation as an important factor in axonal health and survival (Zivraj *et al*., 2010; Gumy *et al*., 2011; Nijssen *et al*., 2018). This in turn indicates that our understanding of local translation and its function needs revision and highlights the necessity for new methods decrypting the axonal transcriptome. Recent advances in stem cell research enable reprogramming of somatic cells into pluripotent stem cells (iPSCs) and their differentiation into different neuronal subtypes. This poses the chance to better understand homeostatic and disease conditions of the brain. Along this line, transcriptome profiling has been used as a valuable tool to investigate changes in the cell body and axonal protrusion of neurons derived from primary embryonic mouse motor neurons and primary embryonic mouse dorsal root ganglia (DRG) (Minis *et al*., 2014; Briese *et al*., 2016). However, these datasets contained high numbers of proliferative and glial marker sets, suggesting contamination by other cellular components or non-neuronal cells (Minis *et al*., 2014; Briese *et al*., 2016). More recently, Nijssen *et al*. developed an approach using a microfluidic device to clearly separate the axon from the cell body with sensitivity similar to single-cell sequencing (Nijssen *et al*., 2018), making it possible to closely examine the axonal transcriptome in healthy and diseased conditions.

To investigate transcripts and pathways critical for axonal maintenance and degeneration, we investigated kinesin family member 1 C (*KIF1C*) dependent changes to the axonal transcriptome. Mutations in *KIF1C*, a motor protein that is causative of an autosomal recessive form of HSP (SPG58, #611302), lead to axonal degeneration of central motor neurons (MN). *KIF1C* is implicated in many functions, including maintenance of Golgi morphology, cell migration, formation of podosomes in vascular smooth muscle cells and macrophages, and MHC presentation on the cell (Kopp *et al*., 2006; del Rio *et al*., 2012; Simpson *et al*., 2012; Theisen *et al*., 2012; Bhuwania *et al*., 2014; Efimova *et al*., 2014; Lee *et al*., 2015). Interestingly, *KIF1C* has also been implicated in long range directional transport of APC-dependent mRNAs and RNA-dependent transport of the exon junction complex (EJC) into neurites, suggesting *KIF1C* can bind RNA in a direct or indirect manner (Pichon *et al*., 2021; Nagel *et al*., 2022; Norris & Mendell, 2023). However, which impact loss of *KIF1C* has on the axonal transcriptome remains elusive.

Using microfluidic chambers, we here provide evidence that RNA-sequencing (RNA-seq) of axons of iPSC-derived cortical neurons (CNs) delivers high-quality, pure, and reliable axonal transcriptome profiles. In agreement with previous studies the axonal compartment contained only a subpopulation of genes compared to the soma and markers for glial and proliferative cells were absent. Importantly, we find that axons not only present with a distinct transcriptome that is clearly separated from the soma but also detected the presence of a set of RNAs coding for transcription factors (TF), previously unknown to be present in the axon.

We show that loss of *KIF1C* leads to widespread changes in this composition and unravel transcriptomic changes that may be relevant for other neurodegenerative diseases. Therefore, this study increases our understanding of homeostatic and disease-related transcriptome conditions in the soma and axons of iPSC-derived CNs.

## Results

### Microfluidic devices enable RNA-sequencing of iPSC-derived human CN axons

To investigate changes in the axonal in relation to the somatic transcriptome in a human neuronal model we performed RNA-seq on axonal and soma samples of iPSC-derived neurons that show characteristics of glutamatergic layer V and VI cortical neurons (CNs). To demonstrate the purity of our iPSC-derived CN culture and ensure expression of cortical markers, immunocytochemical staining and RT-qPCR was performed (Supplementary Figure S1, S2). CN cultures formed complex networks and expressed neuronal markers (β-III-tubulin, TAU), a dendritic marker (MAP2), and markers of cortical layer V (CTIP2) and VI (TBR1) (Supplementary Figure S1A – C). Staining for the astrocytic marker GFAP was negative, indicating high purity of our culture (Supplementary Figure S1D). No morphological differences were noted between wild-type (WT) and *KIF1C-*knockout (KO) CNs. RT-qPCR analysis revealed that iPSC-derived CNs expressed the cortical layer markers FoxG1 and PAX6, the dendritic marker MAP2 and microtubule-associated marker DXC, while undifferentiated iPSCs were negative for these markers (Supplementary Figure S2). Therefore, we conclude that a homogenous culture of neurons was generated that can be used as a cortical *in vitro* cell model. For RNA-seq, CNs were plated into microfluidic devices and the axons were recruited to the empty chamber by using a gradient of glial cell line-derived neurotrophic factor (GDNF), brain-derived neurotrophic factor (BDNF), and nerve growth factor (NGF) (Supplementary Figure S3A, B). Subsequently, each compartment was individually lysed and subjected to RNA-seq (Supplementary Figure S3A; see Methods).

On average, each sample yielded 34.32 million pair end reads, with a mean read length of 109 basepairs (bp). Quality assessments via MultiQC indicated that over 92% of these reads had a Phred score above 30 (Q30), pointing to high sequencing fidelity. We aligned these fastq reads to reference the genome using STAR, achieving an average mapping rate of 89.1%. Following alignment, gene expression was measured in transcripts per million (TPM) values via featureCounts.

In total, 20,199 genes were detectable across all datasets, with a TPM > 1 in at least three RNA samples. Among the whole dataset, the soma samples exhibited an average of 16,159 ± 880 detectable genes, whereas the axon samples demonstrated a significantly lower count, with an average of 5,139 ± 1,642 genes (Figure 1A). An initial principal component analysis (PCA) highlighted sequencing batch effects, which were corrected using the ComBat method from the preprocessCore package. Three axon RNA-seq samples (two WT and one *KIF1C*-KO sample) were removed due to their poor correlation (r < 0.5) with other samples. The remaining samples passing the final QC, resulted in 13 soma (WT: 6; *KIF1C-KO*: 7) and 10 axonal samples (WT: 5; *KIF1C-KO*: 5) (Supplementary Figure S4; Supplementary Table S1).

**Figure 1:**
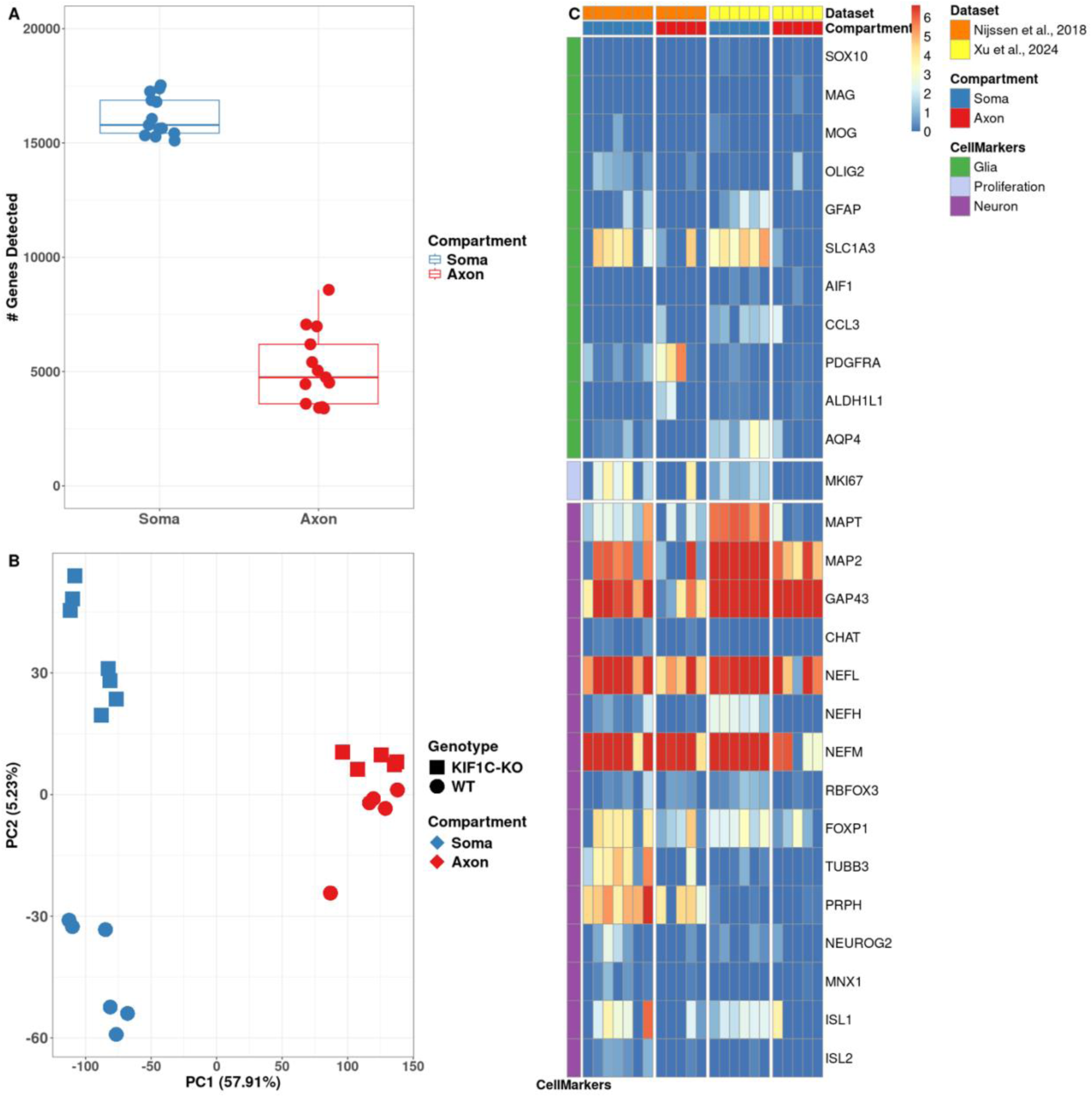
Quality control reveals RNA-seq for axonal compartments is highly specific. **A.** Median number of gene counts detected in the soma (blue) and axonal compartment (red) (TPM>1 in at least 3 samples; across all samples). The number of detected genes in the soma compartment (16,159 ± 880) is approximately three folds higher than in the axon compartment (5,139 ± 1,642). Data is presented as medium and interquartile range. **B.** Post QC PCA scatter plot. PC1 distinguishes the soma (blue) and axonal (red) compartment. PC2 distinguishes WT (circle) and *KIF1C-KO* (square) samples. **C.** Expression log2(TPM) values of glial (green, top row), proliferative (lavender, middle row) and neuronal marker genes (purple, bottom row) in soma (blue, left columns) and axonal compartments (red, right columns), in Nijssen *et al.,* 2018 (orange, left) and our dataset (Xu *et al*., 2024; yellow, right). As in the samples from Nijssen *et al*., our samples show low expression of glial and proliferative marker genes while neural marker genes are highly expressed.

The final PCA after batch correction and sample QC illustrated distinct clustering (Figure 1B). Principal component (PC) 1 prominently differentiated the transcriptome of the axon from soma samples, emphasizing their considerable transcriptional disparities. Concurrently, PC2 separated samples by genetic background—WT versus *KIF1C-KO* (Figure 1B).

To confirm our samples’ cellular composition and purity, we referred to a list of recognized glial, neuronal, and proliferative marker genes (Supplementary Table S2) and visualized their expression in a heatmap (Figure 1C). The analysis revealed that our RNA-seq samples, which passed QC, exhibited strong expression of neuronal markers while showing a marked absence of glial or proliferative marker expression (Figure 1C). Comparison of our data to a recently published dataset, focusing on human stem cell-derived spinal motor axons (Nijssen *et al*., 2018), revealed a congruent expression pattern (Figure 1C).

In summary we demonstrate that our experimental paradigm, that uses microfluidic devices to separate neuronal axons from soma, enables RNA-seq of axonal fractions with a high specificity. Intensive QC revealed high purity of both compartments without contamination by either glial components or cell somas in the axonal compartment.

### Axonal compartments of iPSC-derived CNs show a unique transcriptional RNA profile

Next, we aimed to delineate the transcriptomic differences between soma and axonal compartments. For this, we analyzed 5 WT axon and 6 WT soma samples. On average, we detected 16,207 ± 981 expressed genes (TPM>1) in WT soma samples and 5,075 ± 2080 genes (TPM>1) in axon samples. Of note, we detected a slightly higher number of genes in both compartments compared to two recently published datasets, investigating human-derived lower motor neurons or i3 neurons (Supplementary Figure S5) (Nijssen *et al*., 2018; De Pace *et al*., 2024), using similar cultivation techniques. In contrast, a significantly higher number of genes detected in the axonal compartment was described by a paper using different cultivation, lysis and sequencing techniques (Maciel *et al*., 2018). Comparing genes detected in the axonal compartment of human-derived CNs to previously published sequencing data of mouse-derived motor neurons (Briese *et al*., 2016; Nijssen *et al*., 2018), we only detected little overlap between the datasets (Supplementary Figure S6), emphasizing the need for human-derived cell models in translational research. Interestingly, the analysis of the top 100 expressed genes within each compartment demonstrated that 57 genes exhibited compartment-specific expression and the profile of highly expressed genes varied between compartments: in the soma we predominantly detected protein-coding mRNAs, with diverse functional implications, whereas axonal expression was characterized by a broader spectrum of gene classes, including a notable prevalence of RNAs encoding ribosomal proteins (RP genes) and mitochondrially encoded RNAs (mt-RNA) (Supplementary Figure S7A – C). Of note, 43 of the top overall 100 genes were concurrently expressed in both axon and soma, demonstrating a significant convergence (Supplementary Figure S4B). Among these, two genes encoding microtubule-associated proteins, Stathmin (*STMN*)*1* and *STMN2*, were identified. These proteins are pivotal in axonal development and repair (Rubin & Atweh, 2004; Thornburg-Suresh *et al*., 2023; Lopez-Erauskin *et al*., 2024). Furthermore, we detected *GAP43*, which plays a significant role in axonal growth, particularly during development and regeneration (Denny, 2006; Chung *et al*., 2020).

Genes with expression levels of TPM>1 in a minimum of 3 samples in axon and soma were considered for studying transcriptomic differences. Differential expression analysis identified 14,056 genes differentially expressed (FDR adjusted *p*-value<0.05) between the compartments: 13,745 genes were enriched in the soma compared to the axon and 311 were enriched in the axon compared to the soma (Supplementary Figure S8). PCA, considering all expressed genes in WT compartments, distinctly separated the axon from the soma, with PC1 accounting for 68.3% of the variance (Figure 1B), underscoring the distinct transcriptomic profile of the axonal compartment. Interestingly, the most significantly enriched genes in the axon, displaying a logFC>2 and an adjusted *p*-value<0.05, were involved in ribosomal subunits (*e.g*., *RPL12, RPL39, RPL31*), respiratory chain complex (*e.g*., *MT-ND1, MT-CO1*), ion transport (*e.g.*, *BEST1*), and mRNA splicing (*e.g*., *YBX1*) (Figure 2A). Consequently, GO-term analysis of axon-enriched genes primarily spotlighted ribosome process functions and mitochondrial processes (Figure 2B).

**Figure 2:**
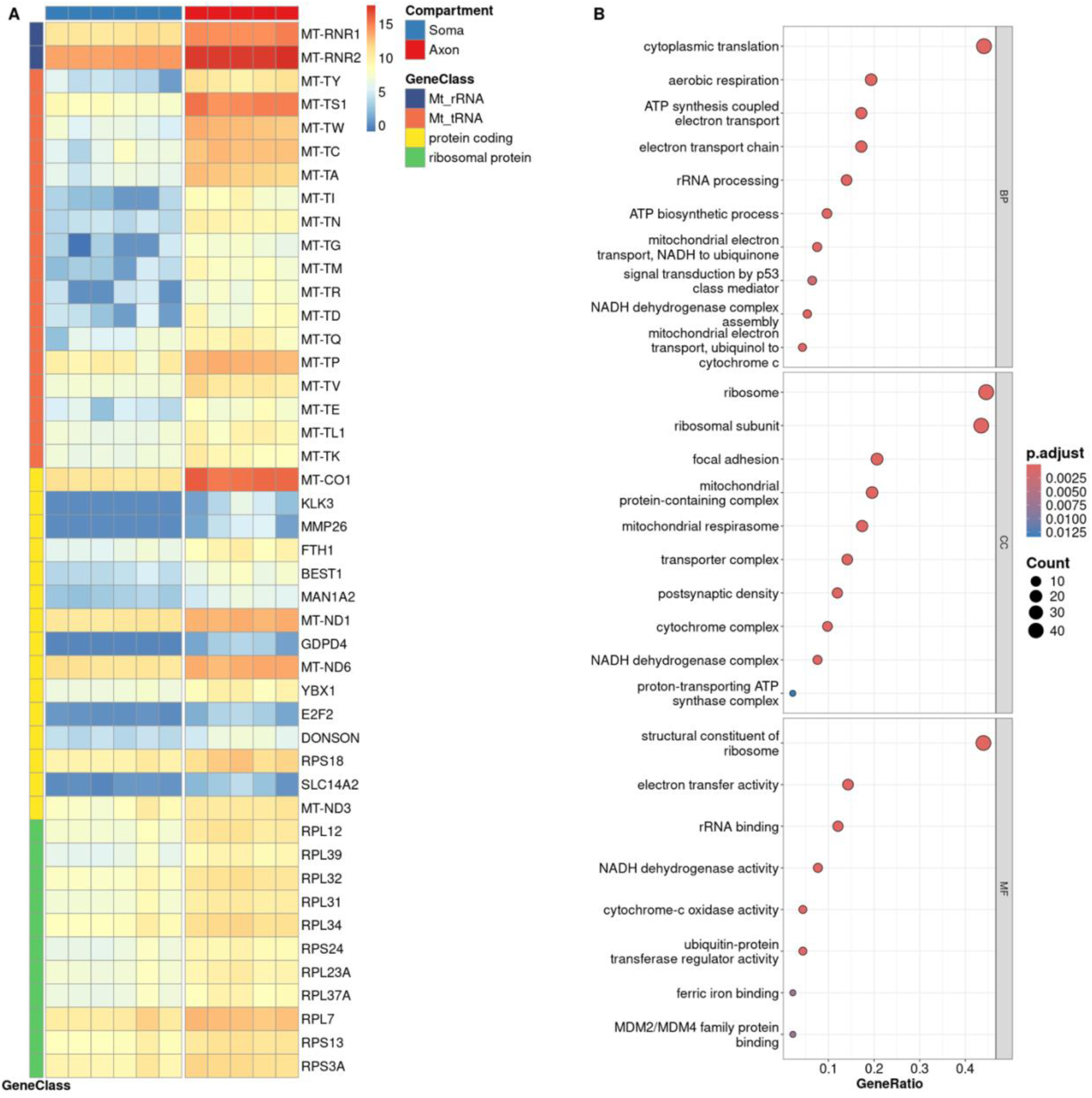
Axonal transcriptome of WT CNs is distinct from the soma transcriptome. **A.** Expression(log2(TPM)) of top genes that are higher expressed in the axonal (red, right column) compared to the soma compartment (blue, left column) (adjusted *p*-value<0.05, logFC>2). Genes are grouped by gene class (purple: mitochondrial (Mt)_rRNA; orange: Mt_tRNA; yellow: protein coding; green: ribosomal protein (RP)). **B.** Gene ontology (GO) term enrichment analysis of axon-enriched genes connected to protein coding or RP genes primarily spotlighted mitochondrial processes and ribosome processes functions. BP: biological process; CC: cellular component; MF: molecular function.

### Axons of iPSC-derived CNs show a unique transcription factor profile

Recent studies have highlighted that transcription factor (TF) mRNAs present in axons can be translated, undergo retrograde transport and alter gene transcription (Cox *et al*., 2008; Wizenmann *et al*., 2009; Ben-Yaakov *et al*., 2012; Ji & Jaffrey, 2012; Guillaud *et al*., 2020; Leboeuf *et al*., 2023). To explore this further, we evaluated expression of a diverse range of TF mRNAs from various organisms in our WT dataset (Shen *et al*., 2023). Upon analyzing the top 50 expressed TF mRNAs separately in axons and somas, we discovered a shared set of 25 TF mRNAs between the two compartments, including key factors like *YBX1*, *SOX4*, *THYN1*, and *SUB1* (Figure 3). In addition to these shared TF mRNAs, each compartment exhibited 25 unique top expressed TF mRNAs, highlighting the specificity of their transcriptional landscapes. Specifically, *GTF3A* and *ATF4* were identified as the leading axonal TF mRNAs, both well-known for their roles in the nucleus initiating transcription (Seifart *et al*., 1989; Anuraga *et al*., 2021) (Figure 3). Moreover, *CREB3* emerged as the most prominent TF mRNA in the soma, playing a crucial role in myelination and axonal growth (Figure 3) (Hasmatali *et al*., 2019; Sampieri *et al*., 2019).

**Figure 3.**
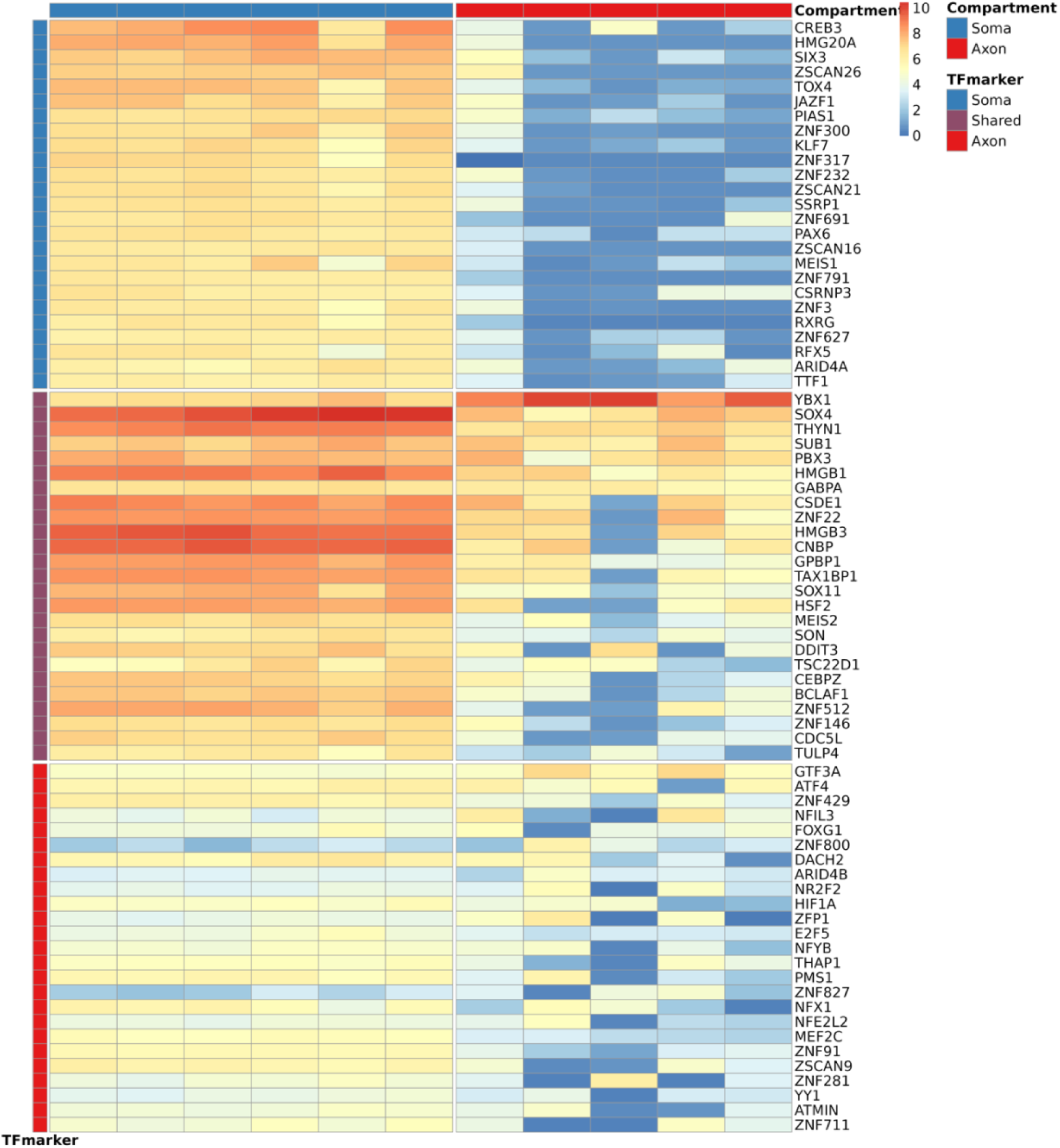
Axon and soma present with specific TF mRNAs. Top expressed transcription factor (TF) mRNAs in the soma compartment (blue, top row), shared (purple, middle row) and axon compartment (red, bottom row), and expression in soma (blue, left column) and axon compartment (red, right column) of iPSC-derived WT CNs. *CREB3* is the top expressed in the soma compartment, while *YBX1*, *SOX4, SUB1* and *THYN1* are shared between axon and soma, and *GTF3A* is top expressed in the axon compartment.

### KIF1C modulates the axonal transcriptome

*KIF1C*-deficiency causes HSP, a disorder associated with prominent length-dependent degeneration of central MN axons. As *KIF1C* has recently been implicated in long-range transport of mRNAs and the EJC (Pichon *et al*., 2021; Nagel *et al*., 2022; Norris & Mendell, 2023), we studied the axonal transcriptome of *KIF1C*-knockout (*KIF1C-KO*) iPSC-derived CNs to highlight transcripts and pathways essential for MN degeneration and maintenance. We did not see gross abnormalities in neuron differentiation, axon growth or axon length in *KIF1C-KO* compared to control CNs. As hypothesized, both axonal and soma compartments in the *KIF1C-KO* cells exhibited markedly reduced *KIF1C* expression when compared with WT compartments (Figure 4A). Looking at the total gene counts, *KIF1C-KO* axons exhibited expression of 5,099 ± 1,642 genes, while WT axon samples displayed 5,075 ± 2080 genes. In the soma we detected 16,207 ± 981 genes in *KIF1C-KO* CNs and 16,118 ± 862 in the WT (Supplementary Table S3). For differential expression analysis, we included genes with a TPM>1 in at least three samples across combined conditions. Ablation of *KIF1C* resulted in a statistically significant differential expression (*p*-value<0.001) of 189 genes within the axon compared to WT axons (Supplementary Figure S9). Intriguingly, this dysregulation was specific to the axon, as only 12 of these also displayed dysregulation in the soma of *KIF1C-KO* CNs (Supplementary Figure S8A, B). *PAX6*, *VIM*, *LGALS1*, *CCND2*, *MIR9-1HG*, *GPR26*, *NR2F2*, *LAMTOR5-AS1* were downregulated in both axon and soma of KIF1C-KO CNs compared to wildtype, and *ZIC2*, *GNRH1*, *GNG8*, *NOL4L* were up regulated in both compartments. 89 genes showed decreased expression in the axon of *KIF1C-KO* CNs, while the remaining 100 exhibited increased expression (Supplementary Figure S9A, B). Furthermore, 21 of the 89 downregulated genes were protein-coding and were totally absent in *KIF1C-KO* CNs (Figure 4C).

**Figure 4.**
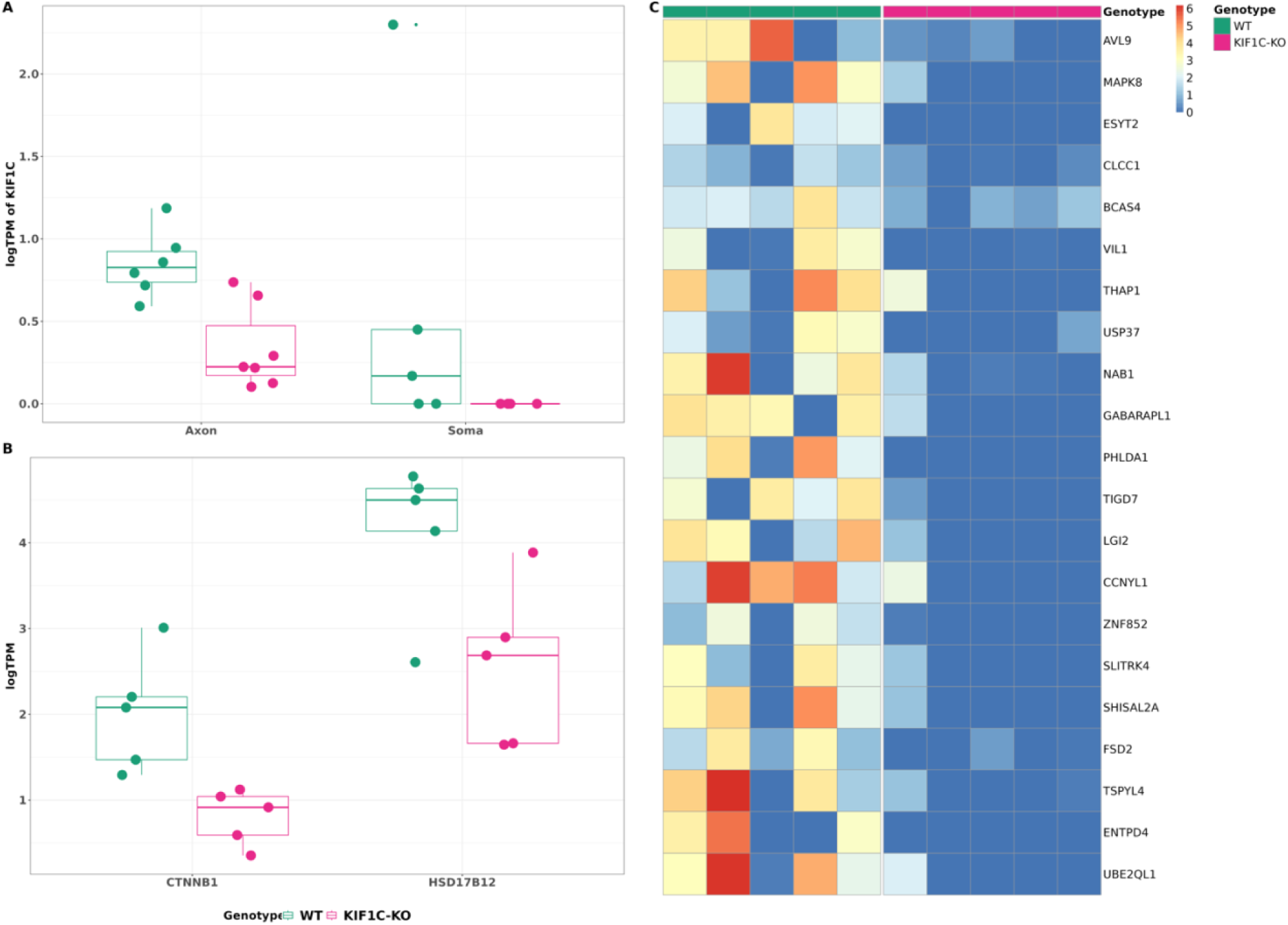
*KIF1C* modulates the axonal transcriptome. **A**. *KIF1C* expression is reduced in the axon and soma compartment of *KIF1C-KO* (magenta) compared to WT CNs (teal). Data is presented as medium and interquartile range. **B.** Two APC target genes show decreased expression in the axons of *KIF1C-KO* compared to WT CNs (WT: teal, *KIF1C-KO*: magenta). Data is presented as medium and interquartile range. **C.** Heatmap of protein-coding that are absent in axons of *KIF1C-KO* compared to WT CNs. All indicated genes are differentially expressed between WT and *KIF1C-KO* axons (*p*-value < 0.001).

Among the transcripts absent or severely downregulated in *KIF1C-KO* CN axons are several key transcripts critical for neuronal and axonal functions (Figure 4C). These transcripts include *ESYT2*, known to play a role in synaptic growth and neurotransmission, *LGI2*, involved in inhibitory synapse assembly (Contreras *et al*., 2019; Kozar-Gillan *et al*., 2023), *NAB1*, a transcriptional repressor with a role in synaptic active zone assembly (Srinivasan *et al*., 2007; Chia *et al*., 2012), *PHLDA1*, involved in injury response (El Soury *et al*., 2018), *CLCC1*, involved in misfolded protein clearance (Jia *et al*., 2015), *THAP1*, implicated in dystonia (DYT6, # 602629) (Cheng *et al*., 2020), and *SLITRK4*, involved in controlling excitatory and inhibitory synapse formation (Yim *et al*., 2013). Additionally, a sub-group of the downregulated genes in the axons of *KIF1C-KO* CNs appear to be involved in axonal pathways. For example, *PPP3CB* is catalogued under KEGG’s axon guidance pathway, while *SLITRK4*, *PAX6*, *MAPK8*, and *CLCN3* are enshrined within the GO term’s axon-related gene set (Table 1).

**Table 1:**
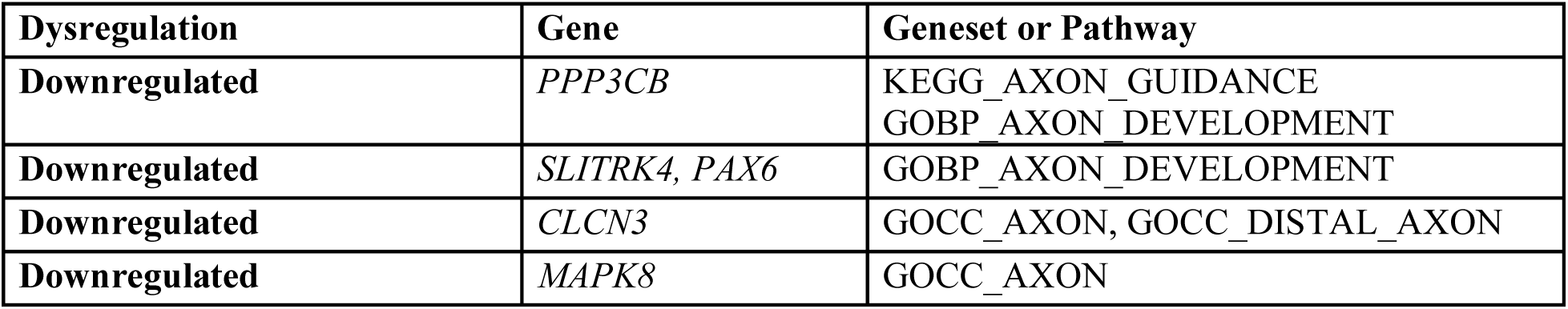
Pathways or gene sets curated for downregulated genes in *KIF1C*-KO axon connected to axon guidance or development.

| Dysregulation | Gene | Geneset or Pathway |
| --- | --- | --- |
| Downregulated | *PPP3CB* | KEGG_AXON_GUIDANCE<br>GOBP_AXON_DEVELOPMENT |
| Downregulated | *SLITRK4*, *PAX6* | GOBP_AXON_DEVELOPMENT |
| Downregulated | *CLCN3* | GOCC_AXON, GOCC_DISTAL_AXON |
| Downregulated | *MAPK8* | GOCC_AXON |

It has been previously described that *KIF1C* plays a role in localizing APC-target mRNAs to cytoplasmic protrusion (Preitner *et al*., 2014). However, only two known targets of APC, *CTNNB1* and *HSD17B12*, were specifically downregulated in axons of KIF1C-KO CNs (Figure 4B).

Taken together, these data show that *KIF1C* plays an important role in the axonal and soma transcriptome and loss of *KIF1C* in CNs leads to widespread transcriptional changes with implications in critical neuronal and axonal functions.

## Discussion

Axonal RNA biology is increasingly implicated in the healthy and diseased CNS. Here, we generated high-quality and pure axonal transcriptomes from iPSC-derived cortical neurons (CNs), using a previously developed RNA-Seq approach (Nijssen *et al*., 2018) with adaptations, and careful application of bioinformatic QC. Our axon samples showed significantly fewer genes than previous datasets from mouse- or human-derived axons (Briese *et al*., 2016; Rotem *et al*., 2017; Maciel *et al*., 2018), which were likely contaminated by non-neuronal cells or somata. Interestingly, our results align with recent, highly specific datasets (Nijssen *et al*., 2018; De Pace *et al*., 2024), though we observed slightly higher gene numbers likely due to our improved lysis protocol, longer paired-end reads, and deeper sequencing. Importantly, our data lacked glial and proliferative markers, indicating highly specific neuronal and axonal cultures. When comparing our data to previously published datasets using mouse-derived cells (Briese *et al*., 2016; Nijssen *et al*., 2018), we found only little overlap of axonal genes, emphasizing the necessity for human-specific research to achieve translational relevance.

Earlier work on human-derived cells focused on axonal transcripts in specific disease contexts, such as ALS in lower motor neurons, (Nijssen *et al*., 2018), and lysosomal-dependent transport in the context of Parkinson’s disease (PD), AD, HD, or ALS using i3 neurons (De Pace *et al*., 2024). In contrast, our work provides a broader overview of molecular profiles in iPSC-derived CNs and explores a previously uncharacterized disease context. Unlike lower motor neurons, residing in the peripheral nervous system, and i3 neurons, which resemble cortical-like neurons but do not fully recapitulate all aspects of CNS-resident neurons, our study focuses on human-derived CNS-resident CNs. Thus, this study offers a broad analysis of axonal and neuronal transcripts relevant to neurodegenerative diseases. Our specific harvesting and sequencing techniques enabled us to identify transcriptional differences impacting general neuronal function and pathology.

Our comprehensive transcriptomic analysis revealed distinct transcriptional landscapes in the axon and soma compartment. While most detected genes specific to the soma compartment represent protein-coding mRNAs, involved in functions essential for neuronal survival and maintenance, axonal compartments showed a more variable range of gene types, including protein coding mRNAs, RNAs encoding RPs, mitochondrial encoded RNAs and lncRNAs. Recent work has provided evidence that local translation of mRNAs in the axons plays an important role in neuronal development, function, plasticity and disease (Li *et al*., 2021). Local translation is regulated by various mechanisms: binding of extrinsic cues to their receptors on the cell surface of axons can trigger local translation (Koppers *et al*., 2019), or chemical signaling through specific cues can activate kinases and phosphorylate RNA-binding-proteins resulting in mRNA translation (Sasaki *et al*., 2010). Interestingly, extrinsic cues like growth factors and nutrients have been shown to activate mTOR signaling which in turn increases translation of axonal mRNAs (Sonenberg & Hinnebusch, 2009; Hornberg & Holt, 2013). Previous studies have shown that RNAs encoding RPs are enriched in mammalian axons, mouse and *Xenopus* growth cones, rat hippocampal dendrites and *Aplysia* neurites (Moccia *et al*., 2003; Poon *et al*., 2006; Zhong *et al*., 2006; Taylor *et al*., 2009; Andreassi *et al*., 2010; Zivraj *et al*., 2010; Gumy *et al*., 2011). Similarly, we detected RNAs encoding for RPs in axons of CNs. While their function remains largely unknown, it is possible that they may be involved in ribosomal repair or renewal of components in already assembled ribosomes (Jung *et al*., 2014). Strengthening this hypothesis, it has been shown that mRNAs encoding RPs are present and locally translated in axons and can be incorporated into axonal ribosomes in a nucleolus-independent manner (Shigeoka *et al*., 2019). Of note, we detected the presence of lncRNAs in the axonal compartment. Recently a lncRNA has been found to be actively involved in local axonal translation (Wei *et al*., 2021), suggesting that lncRNAs in axons have specific functions. The distinct mRNA profile in axons, including nuclear encoded transcripts related to ribosomal subunits, respiratory chain complex, ion transport, and mRNA splicing genes, emphasizes the unique metabolic and synthetic demands of this compartment. This aligns with the hypothesis that axons, despite their dependence on the soma, maintain a degree of autonomy in protein synthesis and energy production, which may be crucial for their function and health.

Transcripts found in both compartments were largely consisting of mitochondrially encoded genes related to NADH metabolism (*e.g*., *MT-ND1,2,4, NDUFA1*), the mitochondrial respiratory chain (*e.g*., *MT-CO1-3, MT-CYB*), or mitochondrial ribosomal functions (*e.g*., *MT-RNR1&2, RPL12*). The rest of the shared signatures were related to cytoskeletal or microtubule organization (*TMSB10, STMN1, STMN2, TUBA1A*), and homeostasis/signaling pathways (*e.g., FTL, GAPDH, DAD1*). These signatures likely reflect the high energy and organizational demands of axon and soma. Interestingly, only one shared gene was related to neurite formation, axon growth, regeneration and plasticity (*GAP43*) (Chung *et al*., 2020). While the role of *GAP43* in the soma is well described, more recently, axonal protein synthesis of *GAP43* was shown to promote axon growth (Donnelly *et al*., 2013; Chung *et al*., 2020).

Recent work has shown that TF mRNAs are not only involved in defining neuronal identity in early development and during survival but function outside the soma. They can be locally synthesized and retrogradely transported back to the nucleus, where they can influence gene transcription and be involved in plasticity, axon pathfinding and neuroprotection (Sgado *et al*., 2006; Cox *et al*., 2008; Sugiyama *et al*., 2008; Wizenmann *et al*., 2009; Torero Ibad *et al*., 2011; Ben-Yaakov *et al*., 2012; Ji & Jaffrey, 2012; Kadkhodaei *et al*., 2013). Therefore, we performed in-depth analysis of enriched TF mRNAs specific to either compartment or common between them. We found that highly expressed TF mRNAs in the soma were related to neural differentiation, morphogenesis, maturation and survival. Of note, two TF mRNAs have been found to be involved in myelination and axonal growth (*RXRG* and *CREB3* (Huang *et al*., 2011; Hasmatali *et al*., 2019; Sampieri *et al*., 2019)), likely reflecting the ongoing growth of axons *in vitro*. In contrast, many TF mRNAs detected in the axonal compartment were related to axon guidance and/or axonal/neuronal regeneration. This is not surprising as several studies have demonstrated that local translation in axons can promote cytoskeletal and membranous growth (Hengst & Jaffrey, 2007; Hengst *et al*., 2009; Gracias *et al*., 2014). We further detected *YBX1* in the axon and soma. *Ybx1* has been found to be enriched specifically in distal motor axons of mice (Nijssen *et al*., 2018). *YBX1* plays an important role in binding and stabilizing cytoplasmic mRNAs, regulating translation and it can mediate anterograde axonal transport (Lyabin *et al*., 2014; Kar *et al*., 2017). Therefore, it is plausible that it can act as an important mediator between axon and soma. Additionally, we found *GTF3A* and *ATF4* enriched in the axonal compartment. These TFs are traditionally linked to transcription initiation and stress response in the nucleus (Seifart *et al*., 1989; Anuraga *et al*., 2021). Importantly, *ATF4*, has previously been shown to be locally synthesized in the axon and retrogradely transported to the soma (Trinh *et al*., 2012; Baleriola *et al*., 2014; Vasudevan *et al*., 2020), suggesting that other TF mRNAs traditionally viewed as nuclear may be, too. In line with this, we detected a substantial number of TF mRNAs with unknown (*e.g*., *ZNF429, ZNF800, ZSCAN9*), or nucleus-related functions, like transcription, cell cycle progression and cell proliferation (*e.g., ATMIN, NFX1, THAP1, E2F5, NR2F2*) in the axon. The discovery of these TF mRNAs signifies a major advancement in our understanding of the molecular mechanisms underpinning axonal functions. These findings not only challenge conventional views of axons as mere passive conduits but also highlight their role as active sites of complex regulatory activities. The potential intra-axonal translation and retrograde transport of translated TFs unveils a dynamic aspect of axons, where they adaptively respond to both internal and external stimuli. This novel paradigm in neuronal biology raises critical questions about the functional roles of TFs within axonal compartments, diverging from their established nuclear functions.

To further investigate changes of the axonal compartment in diseased condition and thus gain an understanding in axonal transcripts essential for axon maintenance, we investigated *KIF1C-KO* CNs. *KIF1C*, a kinesin family member, is implicated in hereditary spastic paraplegia (HSP), a group of heterogenous genetic disorder resulting in degeneration of upper MN axons (Stevanin *et al*., 2008; Caballero Oteyza *et al*., 2014; Dor *et al*., 2014; Novarino *et al*., 2014; Klebe *et al*., 2015; Blackstone, 2018). Indeed, loss of *KIF1C* led to widespread changes of the transcriptome, showing axon-specific effects. This may be explained, as *KIF1C* is the fastest human cargo transporter, and is implicated in long range and highly dynamic transport (Lipka *et al*., 2016). It has long been appreciated that axonal transport is important for survival and health of neurons and disturbance in axonal transport is a key pathological event that contributes to diverse neurodegenerative diseases like AD, polyglutamine diseases, HSPs, ALS, PD, and Charcot-Marie-Tooth (CMT) disease (Perlson *et al*., 2010; Millecamps & Julien, 2013). Additionally, mutations in motor proteins like *KIF5A*, dynein and dynactin have been shown to cause neurodegeneration (Puls *et al*., 2003; Munch *et al*., 2004; Weedon *et al*., 2011; Cady *et al*., 2015; Konno *et al*., 2017), strongly supporting the view that defective axonal transport can directly trigger neurodegeneration. Therefore, spatial dysregulation of transcription due to *KIF1C* loss may uncover transcriptional changes also relevant in other neurodegenerative diseases.

*KIF1C* has previously been shown to transport APC-dependent mRNAs to cell protrusion and it is implicated in the transport of the EJC in neuronal SH-SY5Y cells (Pichon *et al*., 2021; Nagel *et al*., 2022). Therefore, it is not surprising that we see the near complete absence of several key transcripts critical for neuronal and axonal function in axons of *KIF1C-KO* CNs. Interestingly, even though *KIF1C* has been shown to transport mRNAs in an APC-dependent manner, only a subset of the detected missing factors was connected to APC-dependent transport. One explanation for this may be that different motors can be used in different cell types: the transport of *β*-actin mRNA, which accumulates at the leading edge of migrating cells, involves several motors that display cell type and compartment specificity, differing between neurons and fibroblasts (Singer, 1993; Latham *et al*., 2001; Fusco *et al*., 2003; Oleynikov & Singer, 2003; Condeelis & Singer, 2005; Ma *et al*., 2011; Nalavadi *et al*., 2012; Liao *et al*., 2015; Song *et al*., 2015). Interestingly, in mouse fibroblasts, APC-dependent RNAs were involved in pathways functionally relevant to cell movement (Wang *et al*., 2017). However, in this study, genes related to APC-dependent transport did not display functional relevance for movement. Instead, most downregulated RNAs in *KIF1C-KO* axons were related to neurotransmission, the synaptic zone, and misfolded protein clearance. Since the previously mentioned investigations regarding *KIF1C* have been performed in other cell types (Wang *et al*., 2017; Pichon *et al*., 2021; Nagel *et al*., 2022; Norris & Mendell, 2023), it is possible that *KIF1C*-dependent transport in CNs uses different motors, independent of APC. However, by which mechanism transport in CNs is conducted needs to be subject of future investigations. Our study has certain limitations, notably the absence of validation of key transcripts in *KIF1C*-*KO* CNs. Future research should address these limitations to provide a more comprehensive mechanistic understanding.

Taken together, our findings regarding *KIF1C* provide novel insights into the role of motor proteins in axonal transcriptome modulation. Importantly, genes and TFs dysregulated due to *KIF1C* loss may also be implicated in other neurodegenerative diseases.

## Conclusion

Collectively, our study affirms the transcriptional distinctiveness of axonal and soma compartments in iPSC-derived CNs and sheds light on the intricate molecular machinery governing these compartments. The identification of not only compartment-specific gene expression patterns but also compartment-specific TF mRNAs, and the role of *KIF1C* in modulating these, have significant implications for our understanding of neuronal biology and the pathophysiology of MN diseases. Future research focusing on the functional implications of these transcriptional differences and the potential of targeting specific pathways within axons could pave the way for novel therapeutic approaches in neurodegenerative diseases.

## Methods

### Differentiation of iPSCs to CNs

Human induced pluripotent stem cells (iPSCs) were approved for use by the Institutional Review Boards, University of Tübingen Medical School, Germany (approval number: 423/2019BO1). The *KIF1C-KO* line (homozygous knockout; (Nagel *et al*., 2020)) and isogenic control line were grown in essential 8 (E8) medium on diluted Matrigel® solution (1:60, Corning®) coated 6-well plates at 37°C, 5% CO2 and 100% relative humidity. Medium was changed daily. Cells were differentiated into cortical neurons of layers V and VI. The differentiation followed previously established protocols (Hauser *et al*., 2020; Nagel *et al*., 2020; Schuster *et al*., 2020). Briefly, iPSCs were seeded at a density of 3 x 10^5^ cells/cm^2^ on Matrigel-coated plates (Corning®), in E8 medium supplemented with 10µM SB431542 (Sigma-Aldrich) and 500nM LDN-193189 (Sigma-Aldrich). The cells underwent neural induction over 9 days. Post induction, on Day 9, the cells were split at a 1:3 ratio and then cultured in 3N medium with 20ng/mL FGF-2 for an additional 2 days. From Day 11 to Day 26 after induction (DAI 11-26), cells were maintained in 3N medium, with medium changes occurring bi-daily. For RNA-seq, the cells were transferred to microfluidic chambers (XC950, Xona Microfluidics) for cultivation at Day 26. For immunocytochemistry, the cells were transferred to coverslips in a 24-well plate at Day 26.

### Immunocytochemistry

Characterization of cortical neurons was performed by immunostaining. Neuronal markers (β-III-tubulin and TAU), markers of the cortical layer V (CTIP2) and IV (TBR1) and a dendritic marker (MAP2) were used for characterization (Table 2). Additionally, the astrocyte marker glial fibrillary acidic protein (GFAP) was used to ensure culture purity (Table 2). On DAI37, cortical neurons were washed twice with PBS and fixed using 4% (w/v) paraformaldehyde (PFA) (Merck) for 15 minutes at RT. Following, cells were washed three times with PBS and incubated with 5% BSA in 0.1% Triton X/PBS (PBS-T) or 10% NGS in 0.3% PBS-T for 1h at room temperature (RT) (Table 2). The primary antibodies in blocking solution were applied and incubated at 4 °C overnight. After washing three times with PBS-T the secondary antibody (AF488 (Thermo Fisher Scientific, A11001) or AF568 (Thermo Fisher Scientific, A11004)) diluted in blocking solution, was applied for 1 hour at RT in the dark. For nuclear staining, cells were washed three times with PBS-T and incubated with DAPI (1:10.000; Thermo Fisher Scientific, 62248) for 5 minutes at RT. The coverslips were mounted with fluorescence mounting solution (DAKO; Agilent, S3023) on objective slides. Images were acquired using a Zeiss inverted fluorescence microscope Axio Observer 7.

**Table 2:** Antibodies, dilution, and corresponding blocking solution/dilutant used for characterization of iCNs.

| Primary antibody | Dilution | Blocking |
| --- | --- | --- |
| β-III-Tubulin (Sigma Aldrich, T8660) | 1:750 | 5% BSA/0.1% Triton X/PBS |
| CTIP2 (Abcam, ab18465) | 1:500 | 5% BSA/0.1% Triton X/PBS |
| MAP2 (Proteintech, 17490-1-AP) | 1:250 | 5% BSA/0.1% Triton X/PBS |
| TAU (CST, #46687) | 1:1000 | 5% BSA/0.1% Triton X/PBS |
| GFAP (CST, CSIG 80788T) | 1:200 | 5% BSA/0.1% Triton X/PBS |
| TBR1 (Proteintech, 2093-1-AP) | 1:200 | 10% NGS/0.3% Triton X/PBS |

### RT-qPCR analysis

RNA of CNs was reverse transcribed using the Transcriptor High Fidelity cDNA Synthesis Kit (Roche) following the manufacturer’s instructions. Briefly, 500 ng RNA was reverse transcribed to cDNA by incubating the RNA with random hexamer primers for 10 minutes at 65 °C followed by reverse transcription with the Transcriptor High Fidelity Reverse Transcriptase. Quantitative real-time PCR (RT-qPCR) was performed to characterize CNs. cDNA [1:10] was mixed with Light Cycler 480 SYBR Green I Master (Roche) and the respective qPCR primers [10μM] for the housekeeping gene GAPDH and for the dendritic marker MAP2, cortical markers FoxG1 and Pax, and microtubule-associated marker DCX. Transcription level quantification was performed on a LightCycler 480 with a touch down qPCR program in three technical replicates. Expression level quantification was carried out with the LightCycler 480 software tool ‘relative quantification’ by normalizing C_P_ values to the housekeeping gene. Undifferentiated iPSCs were used as a negative control for cortical markers.

### Culturing of cortical neurons (CNs) in microfluidic devices

Microfluidic devices were set up following the procedure outlined by Nijssen *et al*. (Nijssen *et al*., 2018) with modifications regarding the cultivation and lysis protocol (Supplementary Figure S3). Briefly, both compartments of the device were precoated with XC Pre-Coat (Xona Microfluidics), with one compartment receiving a 1-minute incubation and the other a 5-minute incubation. Following the coating, the chambers were washed twice with phosphate-buffered saline (PBS), coated with Polyornithine, incubated at RT for 1 hour, and subsequently coated with a diluted Matrigel® solution (1:45, Corning®), followed by a 1-hour incubation at 37°C. After two washes with 3N Medium, the compartments were filled with 3N Medium supplemented with Rock inhibitor (RI, 1:1000) and maintained at 37°C until neuron loading. DAI26 neurons were washed with PBS, incubated with Accutase + RI for 20 minutes at 37°C, strained through a 70 µm cell strainer, and suspended in 5,5 ml 3N + RI medium. 2 x 10^5^ neurons were loaded into each microfluidic chamber. The next day, 10µM PD0325901 (Tocris) and 10µM DAPT (Sigma-Aldrich) were added to the soma compartment. Cells were kept in this media for four days with media change every two days, followed by incubation with 3N media. 10ng/ml of NGF, BDNF, and GDNF were added to the axonal compartment to attract axons. Media changes were implemented three times per week, and fluid volumes were adjusted to ensure cross-chamber flow and to establish a trophic factor gradient from axonal to somatic compartments. Of note, the soma compartment contains not only cell bodies but also corresponding axonal fractions. For simplicity purposes we will refer to this compartment as soma compartment throughout this paper.

### Harvesting

At day after induction (DAI) 58, the RNA was extracted (Supplementary Figure S3). DAI 58 was chosen for harvesting, because the axonal chamber was adequately overgrown to ensure a high RNA yield for the sequencing approach. No gross morphological differences between wt and *KIF1C-KO* neurons were observed at this time point. The compartments were washed with PBS. For the axonal compartment, 5µl NEBNext Cell Lysis Buffer (NEB, Massachusetts, U.S.), were added directly into the corresponding microgroove. After 3 minutes, the solution was mixed and snap-frozen on dry ice. The soma compartment was lysed using 50µl RLT buffer (Qiagen, Venlo, Netherlands), collected and snap frozen using dry ice. Soma RNA was purified using the RNeasy Mini Kit (Qiagen, Venlo, Netherlands) following the manufacturer’s instructions.

### RNA Seq library construction

Due to the low amount of RNA available in the axon fraction, lysate of approximatively 500 cell was performed in 5 µl of the NEBNext Cell Lysis Buffer and used in the NEBNext Single Cell/Low Input RNA library Prep Kit Low Input following the protocol for Cell instruction. The library molarity was determined by measuring the library size (approximately 330 bp) using the Fragment Analyzer 5300 and the Fragment Analyzer DNA HS NGS fragment kit (Agilent Technologies) and the library concentration (>2 ng/µl) using Qubit Fluorometric Quantitation and dsDNA High sensitivity assay (Thermo Fisher Scientific). The libraries were denaturated according to the manufacturer’s instructions, diluted to 270 pM and sequenced as paired end 100bp reads on an Illumina NovaSeq 6000 (Illumina). The sequencing aimed to achieve a depth of >15 million clusters per sample.

### Short reads RNA-seq processing and differential gene expression analysis

Our RNA-seq pipeline encompassed read quality control (QC), RNA-seq mapping, and gene quantification. Raw RNA-Seq data were processed using the megSAP pipeline, which includes quality control and adapter removal of fastq files. The reads were then aligned to the reference genome using STAR (version 2.7.10a)(32). Gene and transcript expression quantification was performed using FeatureCount (version N). Transcript and gene abundances were expressed as TPM values.

To identify differentially expressed genes between soma and axon compartments of control lines, we first filtered out lowly expressed genes by keeping only those with at least 1 TPM value in at least 3 samples in each compartment. The filtered TPM matrix was taken as a log2 transformation. Differential expression analysis was performed using the Limma-Voom module. For the enrichment analysis of Gene Ontology (GO) terms, we utilized the ReactomePA package in R (version 4.3). This analysis encompassed the evaluation of significantly enriched pathways within the categories of Biological Processes (BP), Cellular Components (CC), and Molecular Functions (MF). The most significantly enriched pathways were meticulously assessed, and the top distinguished functional pathways were selected for visualization.

To compare our data with previously published datasets, we implemented the following procedures: for the dataset by Nijssen *et al*. (2018), we first converted their counts per million (CPM) values to transcripts per million (TPM) values. Subsequently, we required at least two wild-type (WT) samples for both soma and axon to have TPM values greater than 1. For the dataset by De Pace *et al*. (2024) and Maciel *et al*. (2018), which contained only two and 3 WT soma and axon samples in each dataset, we similarly converted the data to TPM values and included genes with TPM values greater than 1 in both samples.

## Abbreviations

ALS: Amyotrophic lateral sclerosis
AD: Alzheimer’s disease
APC: Adenomatous-polyposis-coli-protein
BDNF: Brain derived neurotrophic factor
bp: Basepairs
BP: Biological process
CC: Cellular component
CMT: Charcot-Marie-Tooth
CN: Cortical neuron
CNS: Central nervous system
DAI: Day after induction
DRG: Dorsal root ganglia
EJC: Exon junction complex
GDNF: Glial cell line-derived neurotrophic factor
GFAP: Glial fibrillary acidic protein
GO: Gene ontology
GWAS: Genome-wide association studies
HSP: Hereditary spastic paraplegia
HD: Huntington disease
IPSC: Induced pluripotent stem cell
KIF1C: Kinesin family member 1 C
KO: knockout
MF: Molecular function
MN: Motor neuron
mt: Mitochondrial
NGF: Nerve growth factor
QC: Quality control
PC: Principal component
PCA: Principal component analysis
PFA: Paraformaldehyde
PD: Parkinson’s disease
RP: Ribosomal protein
RT: Room temperature
SMA: Spinal muscular atrophy
SPG58: Hereditary spastic paraplegia type 58
STMN: Stathmin
TF: Transcription Factor
TPM: Transcripts per million
RNA-seq: RNA-sequencing
WT: Wild-type

## Author’s contribution

Author contribution R.S., S.H., L.S., M.Na., J.X., and M.H. designed the research; S.H., M.K., and N.C. established the axon separation and axon seq technique. J.X., M.H., M.Na., P.P., M.K., and M.No. performed the research; R.S., J.X., M.Na. and M.H. analyzed the data; J.A and N.C. performed the RNA-seq library construction. R.S., M.H., and J.X. wrote the paper with input from all authors.

## Acknowledgment and Funding

This work was funded by the Bundesministerium für Bildung und Forschung (BMBF) through funding for the TreatHSP network (grant 01GM1905A and 01GM2209A to RS, grant 01GM2209F to LS and SH), the National Institute of Neurological Disorders and Stroke (NINDS) / National Institutes of Health (NIH) under Award Number R01NS072248 (grant to RS), and the Clinician Scientist Programme PRECISE.net funded by the Else Kröner-Fresenius-Stiftung (grant to RS). RS and LS are members of the European Reference Network for Rare Neurological Diseases – Project ID 739510. NGS sequencing methods were performed with the support of the DFG-funded NGS Competence Center Tübingen (INST 37/1049-1).

We would like to thank Jessica Cielenga for her technical support.

