## Supplementary data for "Unraveling Axonal Transcriptional Landscapes: Insights from iPSC-Derived Cortical Neurons and Implications for Motor Neuron Degeneration"

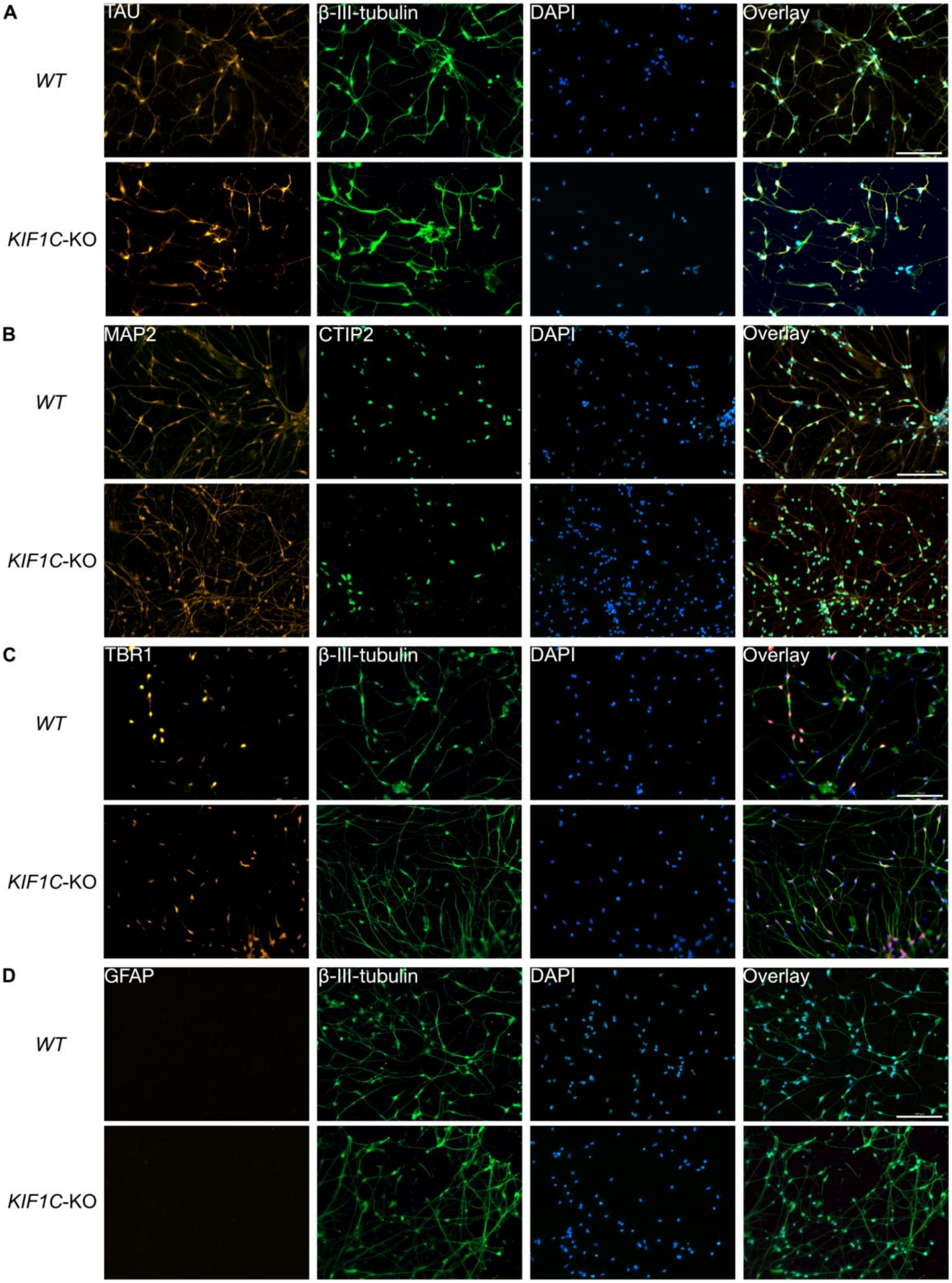

**Figure S1:** **A**. iPSC-derived CNs were stained for axonal marker TAU (red) and neuron-specific marker β-III-tubulin (green), **B.** dendritic marker MAP2 (red) and cortical layer V marker CTIP2 (green), **C** cortical layer VI marker TBR1 (red) and **D** astrocytic marker GFAP (red) in WT (top) and *KIF1C-KO* iPSC-derived CNs (bottom). Nuclei are stained using DAPI (blue). Scale bars represent 100 µm.

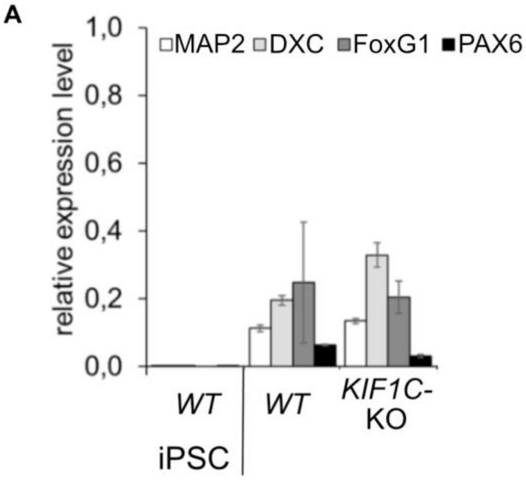


**Figure S2:** **A.** Relative expression levels of dendritic marker MAP2, the microtubule-associated marker DXC, and cortical markers FoxG1, and PAX6 normalized to GAPDH in comparison to undifferentiated iPSCs as assessed by RT-qPCR (data presented as mean ± SD). WT iPSCs (1$^{st}$ column) were negative for aforementioned markers. In contrast, WT (2$^{nd}$ column) and *KIF1C-KO* (3$^{rd}$ column) iPSC-derived CNs were positive.

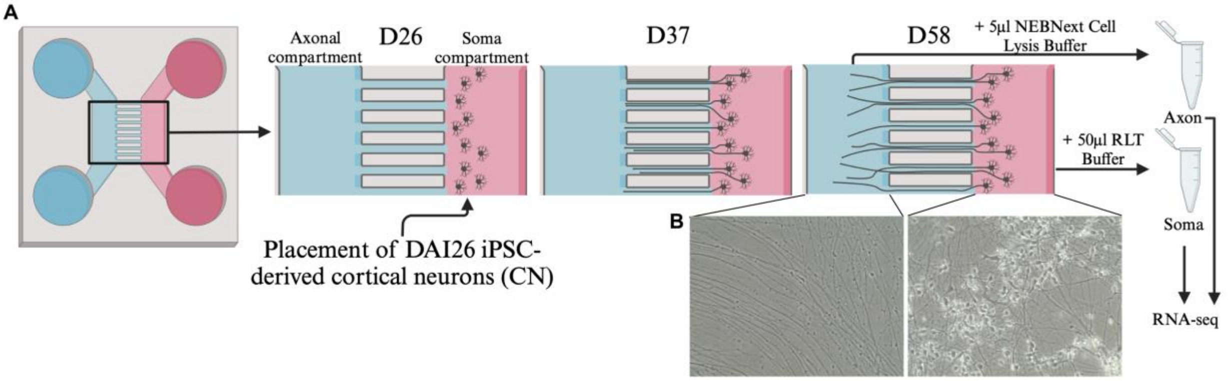


**Figure S3:** **A.** Experimental set-up of microfluidic chamber. iPSC-derived cortical neurons (CN) were placed in the soma compartment (right, pink) of the chamber at day after induction (DAI) 26. Between the chambers a gradient was contained using BDNF, NGF and GDNF to promote axon growth into the axon compartment (left, blue). At DAI58 (D58) the axon compartment was harvested using 5µl NEBNext Cell Lysis Buffer (NEB, Massachusetts, U.S) and the soma compartment was harvested using 50µl RLT Buffer (Qiagen, Venlo, Netherlands). Following, they were sequenced using RNA-sequencing (RNA-seq). **B.**

Representative light microscopic images of the axonal (left) and soma compartment (right). Note the absence of soma in the axonal compartment.

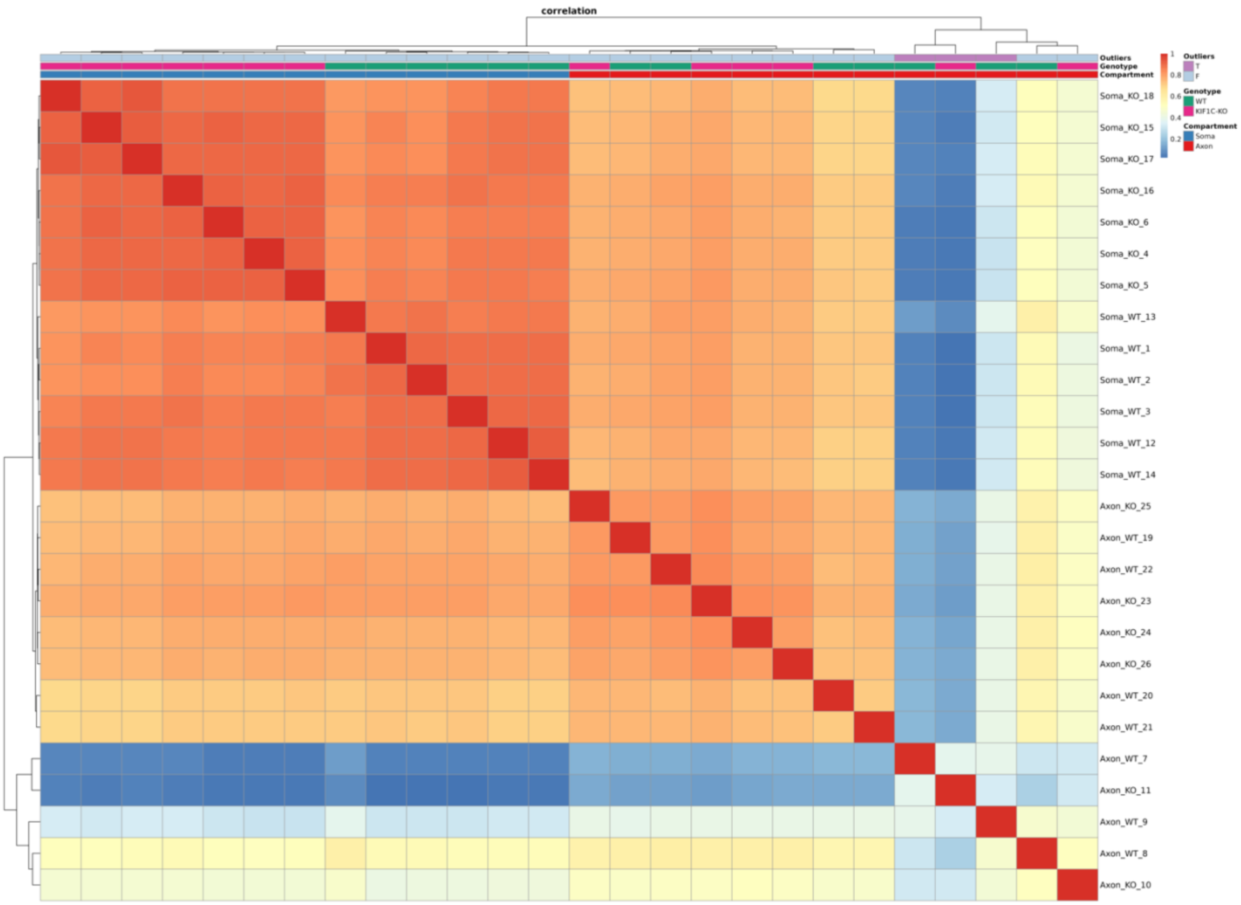


**Figure S4:** Spearman correlation of RNA-seq samples after removal of batch effects. Three samples are considered outliers since they are not correlated to any other samples with a coefficient greater than 0.5. (T= true, F= false; soma: blue; axon: red).

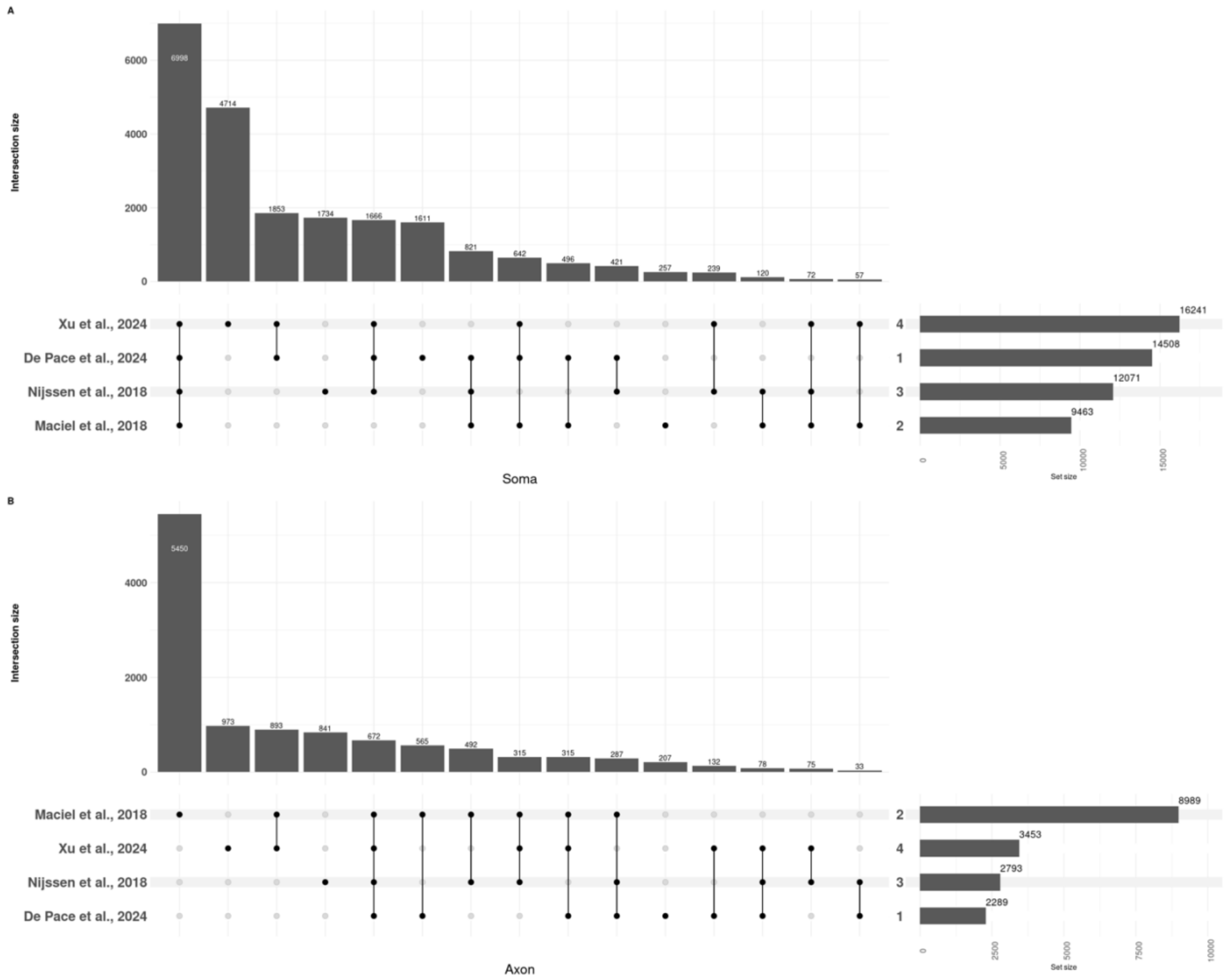


**Figure S5: A.** Upset plot of differential gene expression shows overlapping genes in human-derived cells between De Pace *et al*., (2024; 2nd row), Nijssen *et al.*, (2018, 3rd row), and Maciel *et al*. (2018, bottom row) compared to our study (Xu *et al*., 2024; top row) in the soma compartment. Overall, we detected a higher number of genes in the soma of iPSC-derived CNs (16.241) compared to the numbers reported by DePace *et al*. (14.528) in i3 neurons, and Nijssen *et al* (12.418), and Maciel *et al*. (9.463) in iPSC-derived lower motor neurons. **B**. Overlapping genes between Maciel *et al*., (2018; top row), Nijssen *et al.*, (2018, 3rd row), and De Pace *et al*. (2024, bottom row) compared to our study (Xu *et al*., 2nd row) in the axonal compartment of human-derived cells. We detected similar numbers in the axonal compartment compared to De Pace *et al.*, and Nijssen *et al*.. Maciel *et al.*, detected a significantly higher number of genes in the axon compared to all other studies, that shows a near overlap with detected genes in the soma compartment, implicating soma contamination. The bar plot denotes intersection size, circles denote which comparisons have overlap, and the set size reflects the total number of genes

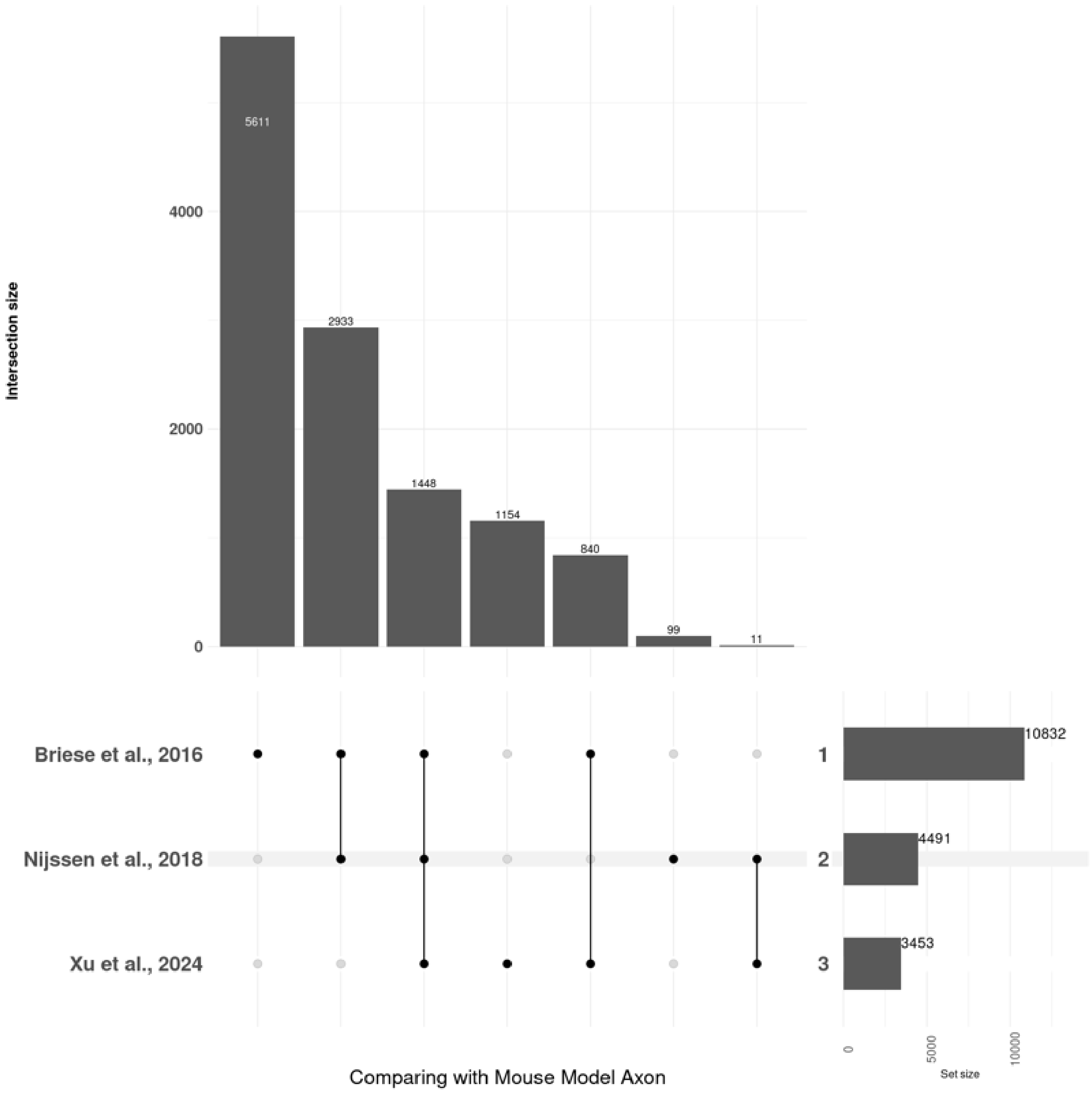


**Figure S6:** Upset plot of differential gene expression shows overlapping genes between human-derived CNs (our data: Xu *et al.*; bottom row) compared to mouse-derived lower motor neurons (Briese *et al.*, (2016; top row); Nijssen *et al.*, (2018; 2$^{nd}$ row)) in the axonal compartment. Overall, we detected fewer transcripts and only little overlap between human- and mouse-derived data. The bar plot denotes intersection size, circles denote which comparisons have overlap, and the set size reflects the total number of genes

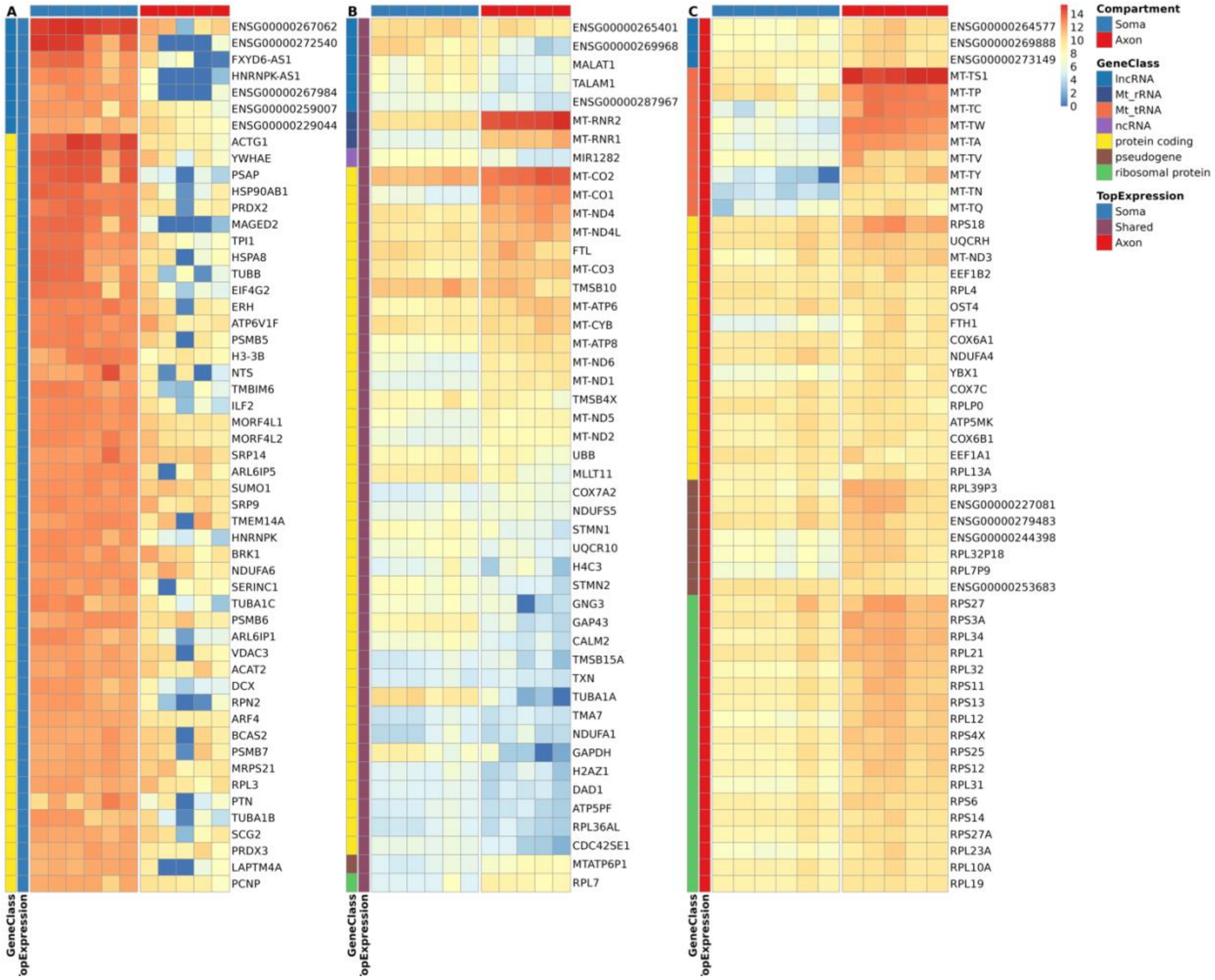


**Figure S7:** Top 100 expressed genes by distribution pattern: **A** soma-specific, **B** common between axon and soma and **C** axon-specific. (Soma, left column: blue; axon, right column: red; lncRNA: indigo; Mt_rRNA: dark blue; Mt_tRNA: orange; ncRNA: lavender; protein coding: yellow; pseudogene: brown; ribosomal protein: green)

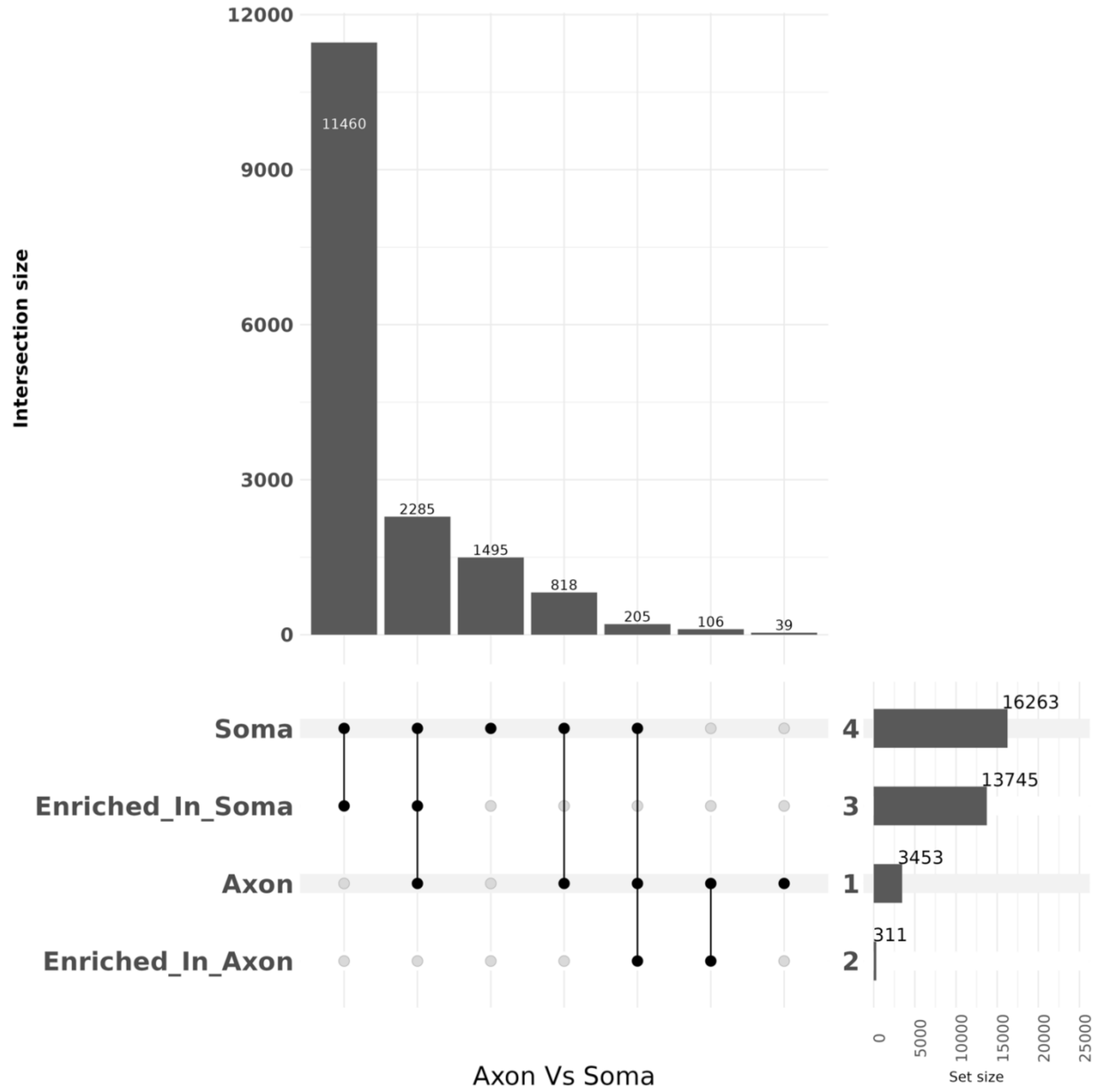


**Figure S8**: Upset plot of differential gene expression between axon and soma compartments of WT CNs. Overall, 16,263 genes were expressed in the soma compartment, of which 13,745 were enriched compared to the axonal compartment. 3,453 genes were expressed in the axonal compartment, of which 311 were enriched compared to the soma compartment. The bar plot denotes intersection size, circles denote which comparisons have overlap, and the set size reflects the total number of genes

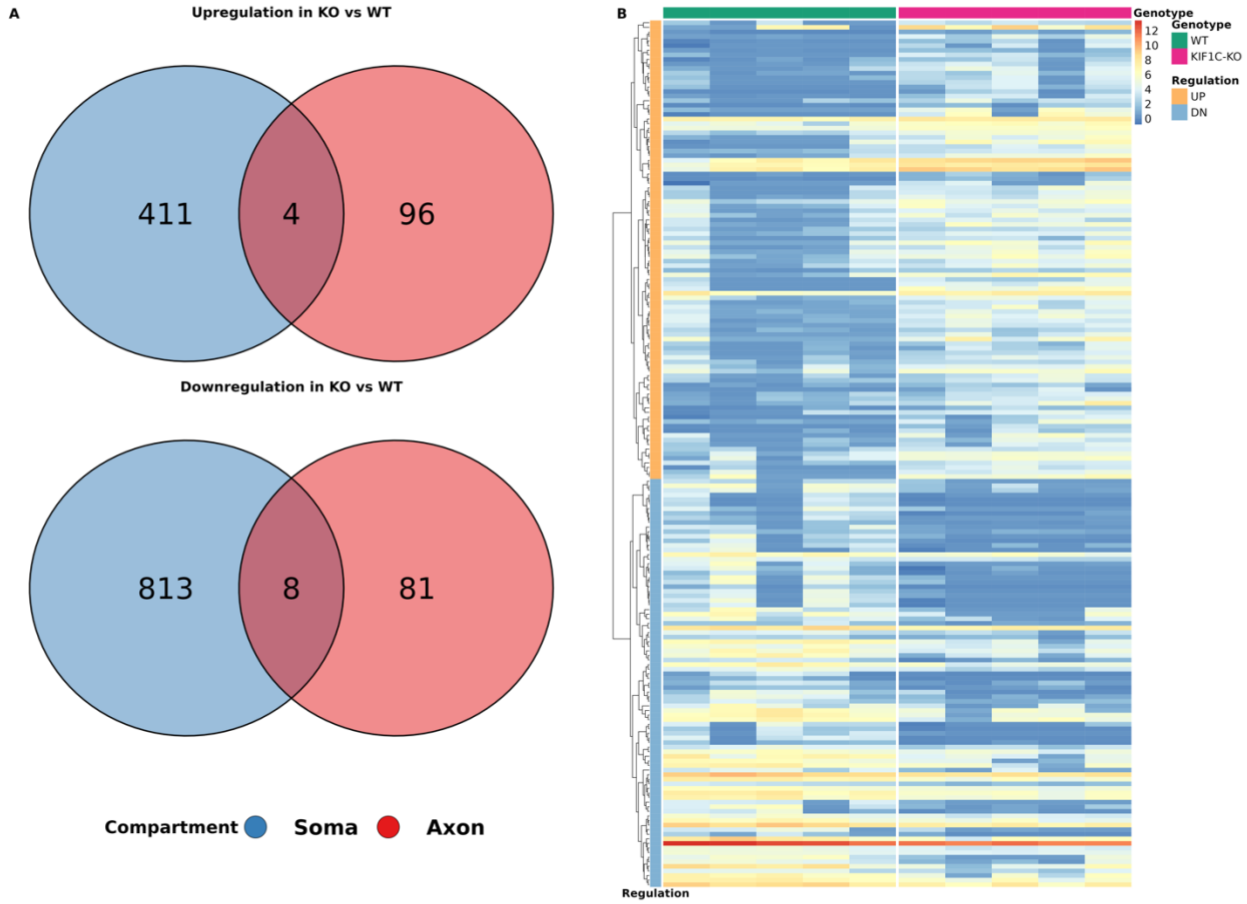


**Figure S9**: **A.** 511 genes are upregulated in *KIF1C-KO vs* WT CNs (upper part; 411 genes upregulated in soma (blue), 96 genes upregulated in axon (red), 4 genes upregulated in both compartments (overlap)). 902 genes are downregulated in *KIF1C-KO* vs WT CNs (lower part; 813 in soma (blue), 81 in axon (red), 8 in both compartments (overlap)). **B.** Differentially expressed genes in the axon of *KIF1C-KO* (right, pink) vs WT CNs (teal, left). 89 genes showed decreased expression (yellow; logFC<0 and p-value <0.001), while the remaining 100 exhibited increased expression (light blue; logFC>0 and p-value <0.001).

| **SampleName Sample ID** | **Type Compartment** | **Cell line** | **CN differentiation batch** | **Production batch Sequencing batch** | **outliers** |
|---|---|---|---|---|---|
| Soma_WT_1 | Soma | CN_control | group4 | Batch1 | F |
| Soma_WT_2 | Soma | CN_control | group4 | Batch1 | F |
| Soma_WT_3 | Soma | CN_control | group4 | Batch1 | F |
| Soma_WT_12 | Soma | CN_control | group6 | Batch2 | F |
| Soma_WT_13 | Soma | CN_control | group6 | Batch2 | F |
| Soma_WT_14 | Soma | CN_control | group6 | Batch2 | F |
| Axon_WT_7 | Axon | CN_control | group4 | Batch1 | T |
| Axon_WT_8 | Axon | CN_control | group4 | Batch1 | F |
| Axon_WT_9 | Axon | CN_control | group4 | Batch1 | T |
| Axon_WT_19 | Axon | CN_control | group6 | Batch2 | F |
| Axon_WT_20 | Axon | CN_control | group6 | Batch2 | F |
| Axon_WT_21 | Axon | CN_control | group6 | Batch2 | F |
| Axon_WT_22 | Axon | CN_control | group6 | Batch2 | F |
| Soma_KO_4 | Soma | CN_KIF1C_KO | group5 | Batch1 | F |
| Soma_KO_5 | Soma | CN_KIF1C_KO | group5 | Batch1 | F |
| Soma_KO_6 | Soma | CN_KIF1C_KO | group5 | Batch1 | F |
| Soma_KO_15 | Soma | CN_KIF1C_KO | group7 | Batch2 | F |
| Soma_KO_16 | Soma | CN_KIF1C_KO | group7 | Batch2 | F |
| Soma_KO_17 | Soma | CN_KIF1C_KO | group7 | Batch2 | F |
| Soma_KO_18 | Soma | CN_KIF1C_KO | group7 | Batch2 | F |
| Axon_KO_10 | Axon | CN_KIF1C_KO | group5 | Batch1 | F |
| Axon_KO_11 | Axon | CN_KIF1C_KO | group5 | Batch1 | T |
| Axon_KO_23 | Axon | CN_KIF1C_KO | group7 | Batch2 | F |
| Axon_KO_24 | Axon | CN_KIF1C_KO | group7 | Batch2 | F |
| Axon_KO_25 | Axon | CN_KIF1C_KO | group7 | Batch2 | F |
| Axon_KO_26 | Axon | CN_KIF1C_KO | group7 | Batch2 | F |

**Table S1:** Investigated samples. KO= *KIF1C-KO* CNs; WT= WT CNs. Outliers are marked (F= false, T=True)

| | Glial Markers | Neuronal Markers | proliferative marker |
|---|---|---|---|
| Genes | SOX10, MAG, MOG, NG2, OLIG2, GFAP, SLC1A3, AIF1, CCL3, PDGFRA, ALDH1L1, AQP4 | MAPT, MAP2, GAP43, CHAT, NEFL, NEFH, NEFM, RBFOX3, FOXP1, TUBB3, PRPH, VACHT, NEUROG2, MNX1, ISL1, ISL2 | MKI67 |

**Table S2** Glial (left), neuronal gene (middle) and proliferative markers (right) used to examine sample purity. Source from https://www.cellsignal.com/pathways/neuronal-and-glial-cell-markers

| Compartment | genotype | # gene per sample | # samples |
|---|---|---|---|
| Axon | WT | 5,075 ± 2,080 | 5 |
| Axon | *KIF1C-KO* | 5,099 ± 1,642 | 5 |
| Soma | WT | 16,207 ± 981 | 6 |
| Soma | *KIF1C-KO* | 16,118 ± 862 | 7 |

**Table S3**: Number of genes detected in the axonal and soma compartment and for *KIF1C-KO* and WT CNs.
